# Nicotinamide mononucleotide treatment increases NAD+ levels in an iPSC Model of Parkinson’s Disease

**DOI:** 10.1101/2020.05.06.080911

**Authors:** Brett K. Fulleylove-Krause, Samantha L. Sison, Allison D. Ebert

## Abstract

Parkinson’s disease (PD) is a common neurodegenerative disorder caused by the loss of dopaminergic neurons in the substantia nigra that leads to severe motor and non-motor deficits. Although the underlying mechanisms of dopaminergic neuron loss is not entirely clear, increasing evidence suggests mitochondrial malfunction as a key contributor to disease pathogenesis. Recently, we found that human PD patient stem cell-derived dopaminergic neurons exhibit reduced nicotinamide adenine dinucleotide (NAD+) levels, an essential cofactor in mitochondrial function and cellular metabolism. In addition, we found that sirtuins, a group of NAD+-dependent deacetylase enzymes that participate in the regulation of mitochondrial function, energy production, and cell survival, displayed decreased activity in PD dopaminergic neurons, thereby suggesting a potential mechanism for dopaminergic loss in PD. Thus, here we tested whether treatment of PD stem cell-derived dopaminergic neurons with an NAD+ precursor could increase NAD+ levels and improve sirtuin activity.

## Introduction

Parkinson’s disease (PD) is a devastating, progressive neurological disorder characterized by the loss of dopaminergic neurons in the substantia nigra and other brain stem nuclei, resulting in decreased dopamine release in the striatum and dysregulated motor output. As the disease advances, PD patients develop tremors, display decreased fine motor control and suffer from non-motor symptoms such as autonomic dysfunction and dementia. As a result, patients are often severely disabled within 10 years of receiving a diagnosis. With an estimated prevalence of 1% in those over 60 years old [1], PD is the second most common neurodegenerative disease worldwide and the incidence and societal impact is likely to increase with the expanding population [2].

Although it is known that death of dopaminergic neurons is the underlying cause of motor symptoms in PD, the exact biochemical processes driving this cell death remain elusive. Mounting evidence from human, animal and cell-based model systems have implicated mitochondrial dysfunction and increased oxidative stress as major drivers in the disease process [3-6]. These two processes are intimately related as oxidative stress damages mitochondrial proteins and impairs autophagy and mitophagy, resulting in increased oxidative damage [3]. Why dopaminergic neurons are particularly vulnerable to aberrant mitochondrial function in PD is not well understood, but it is thought to be related in part to the high metabolic demands of these neurons, increased oxidative stress related to dopamine synthesis, and higher levels of iron and neuromelanin within the substantia nigra pars compacta [3].

While PD is primarily a sporadic disease, several mutations leading to familial PD have been discovered. The G2019S mutation within the leucine rich repeat kinase (LRRK2) gene is the most common cause of familial PD and is phenotypically similar to sporadic PD [7]. In vivo and in vitro studies using model systems expressing the G2019S mutation have demonstrated increased oxidative stress, altered mitochondrial morphology and trafficking, increased mitochondrial fragmentation, and reduced cellular respiration compared to control [3-6]. Previous studies within our lab utilizing induced pluripotent stem cells (iPSCs) harboring the G2019S mutation have found pronounced mitochondrial and sirtuin dysfunction in differentiated dopaminergic neurons [6]. Sirtuins (SIRT) are a class of nicotinamide adenine dinucleotide (NAD+)-dependent protein and histone deacetylases with important regulatory roles in cellular senescence, mitochondrial bioenergetics, DNA repair, and reactive oxygen species production and removal [8]. SIRTs mediate these processes through interactions with a variety of targets, including p53, tubulin, SOD2, PGC-1alpha, and NF-kB that all play critical roles in basic cellular functions [8]. We previously found that dopaminergic neurons generated from LRRK2 G2019S iPSCs have elevated SIRT protein levels, but decreased SIRT activity compared to healthy dopaminergic neurons [6]. Interestingly, we found that these deficits were more pronounced in LRRK2 G2019S iPSC-derived dopaminergic neurons than in cortical neurons derived from the same LRRK2 G2019S iPSCs [6], suggesting a selective vulnerability of dopaminergic neurons. Moreover, we found that LRRK2 G2019S iPSC-derived dopaminergic neurons had reduced NAD levels compared to control dopaminergic neurons [6], which likely impedes SIRT function in dopaminergic neurons and may be contributing to downstream mitochondrial deficits. Previous studies have found that pharmacologically targeting the NAD+ pathway improved mitochondrial and SIRT function in both in vitro and in vivo models of PD [5, 9]. Therefore, here we tested if addition of the NAD precursor nicotinamide mononucleotide (NMN) could increase NAD+ levels and improve SIRT deacetylase function in LRRK2 G2019S iPSC-derived dopaminergic neurons.

## Methods

### iPSCs and dopaminergic neuron differentiation and treatment

Human iPSCs were obtained from a commercially available cell repository (Coriell Institute), maintained as undifferentiated colonies on Matrigel in E8 growth medium. The method for dopaminergic neuron differentiation was adapted from previously published protocols [10, 11]. Briefly, embryoid bodies were cultured in dopaminergic neuron base medium (2% B27, 1% N2, 0.1% β-mercaptoethanol, 50µg/ml laminin, 200ng/ml ascorbic acid, 10mM Y27632, and 1% antibiotic/antimycotic in 25% DMEM, 25% F12, and 50% neurobasal medium) over 14 days. From day 0-3, base medium was supplemented with 8.0µM CHIR-99021, 0.2µM LDN193189, and 40µM SB431542. From days 2-10, the medium was supplemented with 300ng/ml sonic hedgehog, 100ng/ml FGF8, and 2µM purmorphamine. From days 9-14, 1µM DAPT was added to the medium. From days 11-14, 10µg/ml BDNF, 10µg/ml GDNF, and 1µM db-cAMP were added to the medium. On day 14, EBs were dissociated with TrypLE and plated onto Matrigel-coated 6-well plates (200,000 cells/well) and coverslips (30,000 cells/coverslip in 24-well plates) in maturation medium (base medium plus 10mM DAPT, 10g/ml BDNF, 10µg/ml GDNF, and 1mM db-cAMP). Medium was replaced every other day until the end of the experiment. At 42 days of differentiation, control and LRRK2 neurons were treated with 1mM NMN or maturation medium alone for 24hrs.

### Immunocytochemistry

Cells were fixed in 4% PFA in PBS. Following blocking of nonspecific labeling and permeabilization in 5% donkey serum and 0.2% TX-100 in PBS, cells were incubated in appropriate primary antibody and then the corresponding secondary antibody. Hoechst dye was used to label nuclei. Primary antibodies used were rabbit anti-tyrosine hydroxylase (Pel-Freez, P40101, 1:1000) and mouse anti-beta III tubulin (Sigma, T8660, 1:2000).

### Western Blot

Whole cell lysates were isolated from PD and healthy dopaminergic neurons using 1x Chaps buffer with protease inhibitors. Bradford assay was used to asses protein samples and 20ug of protein per sample was run on 12% precast polyacrylamide gels. Gels were electrophoresed at 105 volts for 90 minutes and transferred to PVDF membrane. Protein content of samples was assessed using REVERT total protein stain (Li-COR, 926-11011) and then probed following standard western blot methods. Primary antibodies used were as follows: rabbit anti-SIRT 1 (Cell Signaling, 9475, 1:1000), rabbit anti-SIRT 2 (Cell Signaling, 12672, 1:1000), rabbit anti-SIRT 3 (Abgent, AP6242a, 1:1000), goat anti-PGC1 alpha (Abcam, ab106814, 1:1000), mouse anti-acetylated tubulin (Sigma, T7451, 1:1000), and rabbit anti-SOD2 (Abcam, ab137037, 1:1000). Appropriate species corresponding red and green tagged secondary antibodies were used. Fluorescence was measured using an Odyssey CLx imager and signal intensity quantified using Image Studio software.

### NAD Assay

Quantification of NAD content in samples was carried out using a colorimetric assay kit (Sigma Aldrich). Samples from a single well (200,000 cells) of each experimental group were treated with NADH/NAD extraction buffer and deproteinized using a 10kDa cut-off spin filter to prevent enzymatic degradation of NADH. A standard curve was generated using serial dilutions of 1mM NADH. Samples were then treated with an NAD cycling enzyme mix in duplicate and incubated for 3 hours. Absorbance was measured at 450nm. Following correction for background, NAD content was calculated using the standard curve.

### Statistical Analysis

A total of five independent dopaminergic differentiations were carried out for data analysis using one control and one homozygous LRRK2 G2019S iPSC line. Data were analyzed using ANOVA with Bonferroni multiple correction using GraphPad Prism software. Results were considered statistically significant at p< 0.05.

## Results

iPSCs from a healthy individual and an unrelated individual with confirmed LRRK2 G2019S PD were differentiated into dopaminergic neurons in vitro using protocols adapted from previously established methods [10, 11]. We have previously found that three independent LRRK2 G2019S iPSC lines all showed consistent dopaminergic neuron differentiation efficiency and a similar reduction in mitochondrial function and SIRT deacetylase activity compared to multiple independent control iPSC lines [6]. Therefore, for this pilot study, we chose to test only a single PD iPSC line and a single control iPSC line. To assess successful differentiation into ventral mesencephalon dopaminergic neurons, we used immunocytochemistry to label cells for Tuj1, a beta-III tubulin specific to neurons, and tyrosine hydroxylase (TH), a dopaminergic neuron specific enzyme responsible for dopamine production. Consistent with our and others’ previous work [6, 10], we observed robust Tuj1 expression with a proportion of the neurons also expressing TH immunofluorescence (Fig. 1).

**Figure 1:**
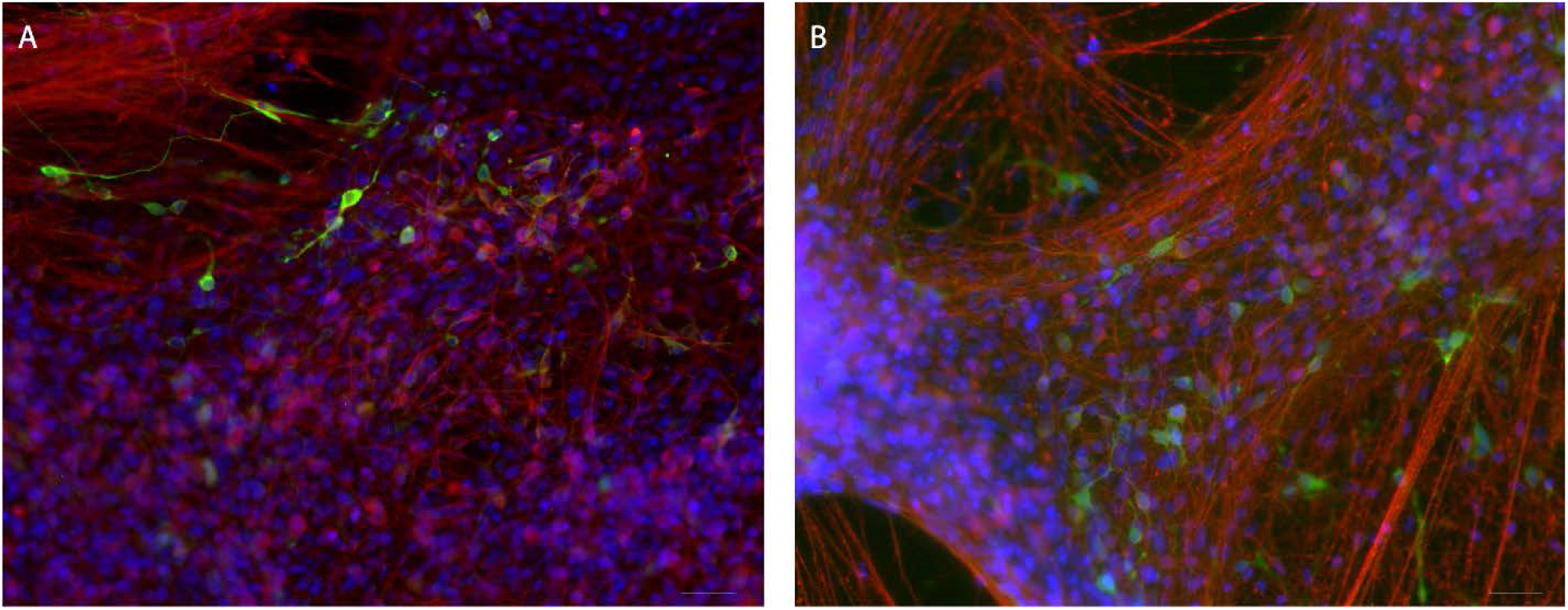
Representative images of dopaminergic neurons generated from healthy (A) and LRRK2 G2019S (B) iPSCs as demonstrated by immunocytochemistry for tyrosine hydroxylase (green) and βIII tubulin (red). Nuclei are labeled with Hoechst nuclear dye (blue). Scale bar = 100µm

Next, we tested whether NMN treatment could effectively increase NAD+ levels in G2019S dopaminergic neurons. We first found that untreated G2019S PD neurons had approximately half of the NAD+ concentration compared to control neurons (Fig 2A). Although this reduction did not reach significance, the trend is consistent with our previous data [6]. NMN treatment increased total NAD+ in G2019S PD neurons compared with untreated G2019S PD neurons to a level comparable to untreated healthy control neurons (Fig. 2A). Considering that NAD+ is required for SIRT function, we next examined the effect of NMN treatment on SIRT deacetylase activity. Similar to our previous data [6], western blot analysis showed LRRK2 G2019S dopaminergic neurons exhibited increased expression of SIRT 1, 2, and 3 (Fig. 2B-D). However, NMN treatment did not dramatically alter overall expression levels of any surtuin tested (Fig 2B-D). We next assessed levels of acetylated targets of SIRT 1, 2, and 3 to determine if NMN treatment also impacted SIRT function. We found no change in expression of acetylated targets PGC1α and SOD2 (data not shown) suggesting that the NMN treatment was insufficient to alter SIRT 1 and 3 activity, respectively. However, treatment with NMN induced a significant 2.5-fold reduction in expression of acetylated α-tubulin in NMN-treated G2019S dopaminergic neurons compared to untreated G2019S neurons (Fig. 2E) indicating a partial restoration of SIRT 2 activity despite no overt change in SIRT 2 levels. Together, these data indicate that NMN treatment may preferentially target and restore SIRT 2 function in LRRK2 G2019S dopaminergic neurons. However, these data also indicate that an increase in NAD+ level is not sufficient to impact overall sirtuin function suggesting that other mechanisms are likely contributing to sirtuin malfunction in G2019S iPSC-derived PD dopaminergic neurons.

**Figure 2:**
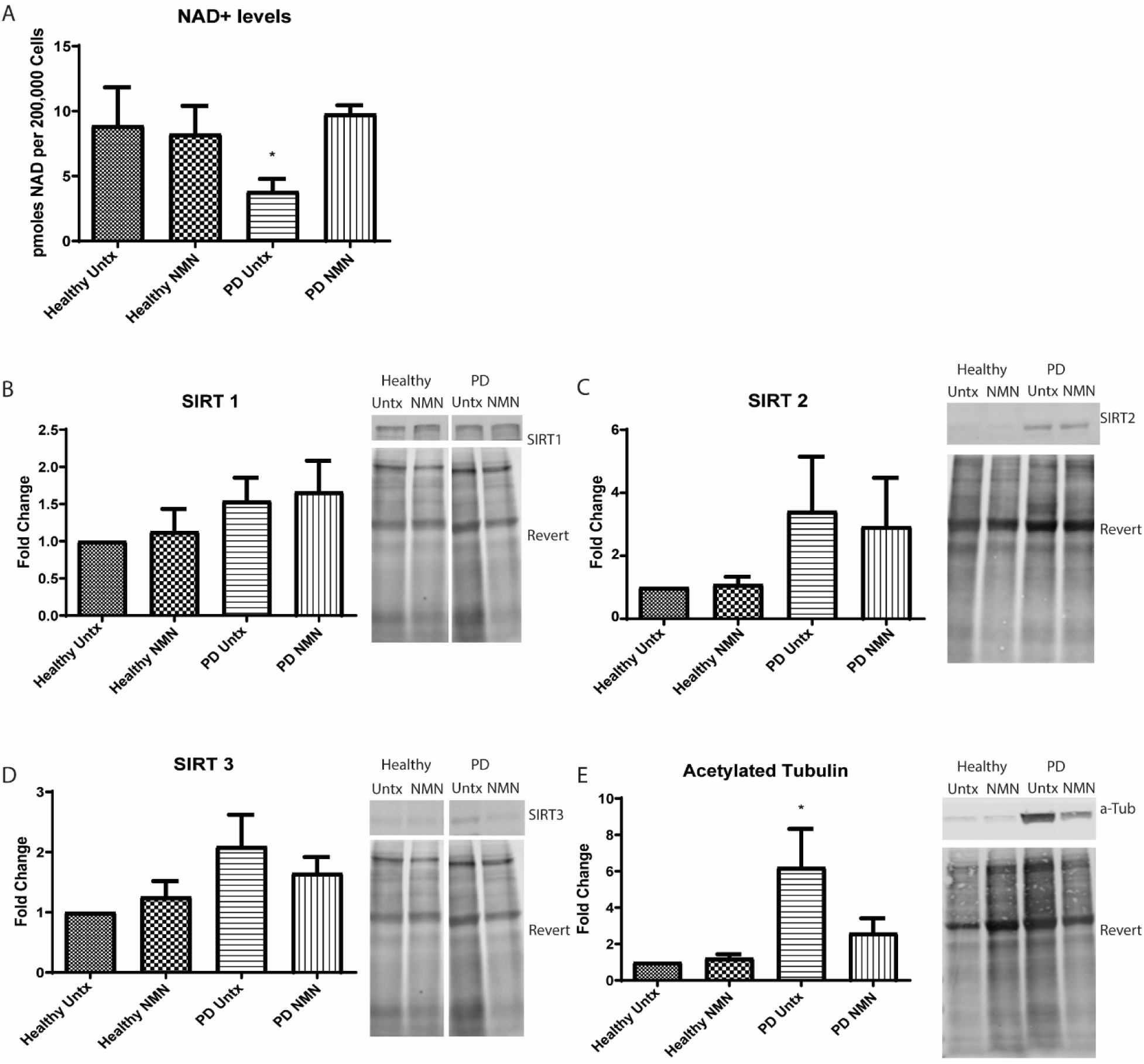
NMN treatment increased NAD+ levels in LRRK2 G2019S iPSC-derived dopaminergic cultures, but it had little impact on SIRT expression or deacetylase activity. (A) NAD+ levels were significantly increased in NMN treated PD iPSC dopaminergic neurons compared to untreated (untx) PD iPSC dopaminergic neurons. NMN treatment had no significant effect on NAD+ levels in healthy dopaminergic neurons. (B-D) Western blot analysis showed no impact of NMN treatment on SIRT 1 (B), SIRT 2 (C), or SIRT 3 (D) protein expression levels in either healthy or PD dopaminergic neurons compared to untreated (untx) dopaminergic neurons. (E) Acetylated α-tubulin (a-tub) was significantly increased in untreated PD dopaminergic neurons, and NMN treatment induced a 2.5-fold reduction in expression of acetylated α-tubulin in PD dopaminergic neurons (E). Protein was normalized to revert total protein stain. *p<0.05 by ANOVA with Bonferroni correction. n=5 independent differentiations. SIRT 1 and SIRT3 were performed on the same membrane, so the revert protein stain is the same in B and D.

## Discussion

Sirtuins are integral in protecting the cell during times of oxidative stress or increased metabolic demand due to their role in the regulation of antioxidant defense, mitochondrial function, and mitochondrial trafficking [8]. As dopaminergic neurons are highly active and selectively damaged in PD, chronic alterations in SIRT function may result in accumulating cellular damage and ultimately cell death with aging in PD patients. In the present study, NMN treatment increased NAD levels, which had the most robust effect on SIRT 2 function in human LRRK2 G2019S iPSC-derived dopaminergic neurons. SIRT 2 is involved in mitochondrial trafficking via deacetylation of α-tubulin. Mitochondrial trafficking from the cell body to dendrites and axons is a critical function that neurons undergo constantly for energy maintenance and survival [12], and dysfunction of this process has been found in a number of neurogenerative diseases, including PD. In this regard, decreased SIRT 2 activity in G2019S DA neurons may be playing a role in the mitochondrial trafficking defects observed in these cells [6]. However, whether NMN treatment improves mitochondrial trafficking in G2019S DA neurons has yet to be tested.

A single administration of 1mM NMN was used based on preliminary data and previous studies showing an effect in iPSC derived neurons from PD patients with mutations in beta-glucocerebrosidase (GBA) [5], but this study did not evaluate the effect on SIRT levels or function. Another study demonstrated that increasing levels of NAMPT, the rate limiting enzyme in the NAD+ salvage pathway, increased SIRT1 activity in 6-hydroxydopamine (6-OHDA) treated PC-12 cells [9]. Although we observed an increase in SIRT2 activity in LRRK2 G2019S iPSC-derived dopaminergic neurons (Fig 2), we did not observe an effect on SIRT1 or SIRT3 expression or function. The discrepancy in results are not entirely clear, but they could be due to differences in model system used (PC-12 cells vs human iPSC-derived dopaminergic neurons), PD stressor (6-OHDA toxicity vs endogenous LRRK2 mutation), or pharmacological treatment (NAMPT vs NMN). Future studies should evaluate the efficacy of different NMN dosage on the sirtuin activity and overall mitochondrial energetics and movement. In addition, limitations of iPSC models must be considered when evaluating pharmacological treatment, including the lack of complex interactions between neuronal, glial, immune, and vascular cell types when working in 2D in vitro cultures. Despite these caveats, iPSCs remain a powerful disease modeling method as they allow for in vitro studies of human cells with a known disease genotype, giving a translational advantage over induced disease models or animal studies. Although additional research is needed, the present study shows that NAD+ precursor treatment can improve overall NAD+ levels in dopaminergic neurons, but the overall impact on SIRT deacetylase activity was modest suggesting that additional modulators may be needed to provide a more robust and therapeutically beneficial effect.

Levodopa administration is the standard treatment for PD patients, but levodopa loses efficacy over time and is associated with significant side effects at higher doses indicating that additional therapeutic options are needed. As we gain greater insight into the genetic and neurobiological events underlying the neuronal death observed in PD, research into disease modifying neuroprotective therapies will advance. The data presented here offer another potential avenue for PD therapeutics. As NAD+ supplements are readily available to the general public, it will be important to further evaluate the effect of NAD+ precursor treatment on neuron health and disease to better understand the potential therapeutic benefits and drawbacks to their use.

## Author contributions

Conception of design: SLS, ADE; Collection of data: BFK, SLS; Analysis of data: BFK, ADE; Supply of resources: ADE; Drafting and editing of manuscript: BFK, SLS, ADE

## Acknowledgements

BFK was supported in part by the National Institute on Aging of the National Institutes of Health (NIH) training grant award (2 T35AG29793). The content is solely the responsibility of the authors and does not necessarily represent the official views of the NIH.

